# Requirements for coregistration accuracy in on-scalp MEG

**DOI:** 10.1101/176545

**Authors:** Rasmus Zetter, Joonas Iivanainen, Matti Stenroos, Lauri Parkkonen

## Abstract

Recent advances in magnetic sensing has made on-scalp magnetoencephalography (MEG) possible. In particular, optically-pumped magnetometers (OPMs) have reached sensitivity levels that enable their use in MEG. In contrast to the SQUID sensors used in current MEG systems, OPMs do not require cryogenic cooling and can thus be placed within millimetres from the head, enabling the construction of sensor arrays that conform to the shape of an individual’s head. To properly estimate the location of neural sources within the brain, one must accurately know the position and orientation of sensors in relation to the head. With the adaptable on-scalp MEG sensor arrays, this coregistration becomes more challenging than in current SQUID-based MEG systems that use rigid sensor arrays.

Here, we used simulations to quantify how accurately one needs to know the position and orientation of sensors in an on-scalp MEG system. The effects that different types of localisation errors have on forward modelling and source estimates obtained by minimum-norm estimation, dipole fitting, and beamforming are detailed.

We found that sensor position errors generally have a larger effect than orientation errors and that these errors affect the localisation accuracy of superficial sources the most. To obtain similar or higher accuracy than with current SQUID-based MEG systems, RMS sensor position and orientation errors should be < 4 mm and < 10°, respectively.

## 1 Introduction

Magnetoencephalography (MEG) is a non-invasive functional neuroimaging method for investigating electric neuronal activity inside the living human brain (Hämäläinen et al 1993; Hansen et al 2010). MEG functions by measuring the magnetic field produced by neural currents in the brain using sensors positioned around the head. The MEG signal is complementary to that of electroencephalography (EEG), in which the potential distribution caused by neural activity is measured using electrodes placed on the scalp.

So far, the magnetometer employed for MEG has almost exclusively been the low-T_c_ superconducting quantum interference device (SQUID). These sensors require a cryogenic temperature that is typically attained by immersing SQUIDs in liquid helium (*T* ≈ 4.2 K ≈ −269°C). The necessary thermal insulation keeps SQUIDs at least 2 cm away from the scalp in most commercial systems and makes the construction of adaptable sensor arrays extremely challenging. Since sensitivity and spatial resolution are related to the distance between the sources and the sensors, the need of cryogenics eventually results in a considerable loss of signal amplitude and spatial resolution (Boto et al 2016; Iivanainen et al 2017). Since the sensor positions cannot be adapted to the head shape of individual subjects, the sensor–scalp-distance is further increased especially when measuring children and infants.

New sensor technologies with sensitivity high enough for MEG have emerged recently; optically-pumped magnetometers (OPMs) (Budker and Romalis 2007; Budker and Kimball 2013) and high-T_c_ SQUIDs (Öisjöen et al 2012) hold promise as alternatives to low-T_c_ SQUIDs. These new sensor types do not require the same degree of thermal insulation and can thus be placed much closer to the scalp, considerably boosting the sensitivity to neural sources. Especially the so-called zero-ñeld OPMs that operate within the spin exchange relaxation-free (SERF) regime (Allred et al 2002) appear promising as they offer high sensitivity while not requiring any cryogenics. Additionally, SERF OPMs can be miniaturised (Shah et al 2007; Knappe et al 2011), enabling the construction of high-density sensor arrays. SERF OPMs with sensitivities better than 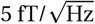 have been demonstrated (Kominis et al 2003; Griffith et al 2010; Colombo et al 2016; Knappe et al 2016).

One could envision EEG-cap-like MEG sensor arrays utilising OPMs, in which the shape of the array conforms to the shape of the head. Recent simulation studies (Boto et al 2016; Iivanainen et al 2017) have shown these *on-scalp* MEG systems to have significantly higher sensitivity to neural sources compared to SQUID-based systems.

To be able to determine where in the brain the measured neuromagnetic signal originates (i.e. to perform source estimation), one needs to accurately know the position of sensors in relation to the head. In practice, source estimation is performed in conjunction with structural magnetic resonance images (MRI), enabling modelling of the head geometry at an individual level. For such modelling to be possible, the MEG data have to be coregistered with the MRI, i.e. the data from both modalities have to be transformed into a common coordinate system.

In current SQUID-based MEG systems, the sensors are rigidly mounted to the insert inside the cryogenic vessel, dewar. Thus, for coregistration, only the position and orientationmjn of the subject’s head in relation to the dewar need to be determined. In an on-scalp MEG system with non-rigid array geometry, coregistration becomes more challenging as each sensor needs to be individually localised with respect to the head. The nature of coregistration error also changes: in SQUID-based MEG systems, the coregistration error is mainly a systematic shift of the whole array, while in on-scalp MEG systems the dominant type of coregistration error may be sensor-wise.

Coregistration for on-scalp MEG systems is similar to that for EEG, but in contrast to EEG, one needs to know the sensor orientation in addition to position. While the coregistration accuracy required for useful source estimation in EEG is agreed to be roughly < 5 mm (Brinkmann et al 1998; Wang and Gotman 2001; Koessler et al 2007), the requirements for on-scalp MEG have so far been mostly unexplored. Recently, Andersen and colleagues (2017) empirically showed the importance of not only determining both the position but also orientation of on-scalp MEG sensors using a single high-T_c_ SQUID.

In this work, we systematically determine how accurately one needs to know the sensor positions and orientations in on-scalp MEG systems for uncompromised forward and inverse modelling. We simulate a hypothetical EEG-cap-like OPM sensor array and investigate the effect of sensor-wise position and orientation error. We quantify the effect of these errors on the forward models as well as the performance of three common source estimation procedures — minimum-norm estimation, dipole modelling and beamforming — in the presence of sensor localisation error.

## 2 Materials and methods

### 2.1 Anatomical models

Existing T1-weighted magnetic resonance images obtained from ten healthy adults (7 males, 3 females) using a 3T MRI scanner were used. The FreeSurfer software package (Dale et al 1999; Fischl et al 1999a; Fischl 2012) was used for pre-processing the MRIs and for segmentation of the cortical surfaces.

For each subject, the surfaces of the skull and scalp were segmented using the watershed approach (Ségonne et al 2004) implemented in FreeSurfer and MNE software (Gramfort et al 2014). These surfaces were thereafter decimated to obtain three boundary element meshes (2 562 vertices per mesh). The neural activity was modelled as a primary current distribution constrained to the surface separating the cortical gray and white matter and discretised into a set of 10 242 current dipoles per hemisphere. For dipole modelling simulations, a sparser nonoverlapping discretisation with 20-mm inter-source spacing was constructed. For minimum-norm estimation, sources were assumed to be normal to the local cortical surface, while for dipole modelling their orientation was not constrained. For visualisation and group-level statistics, individual brains were mapped to the average brain of the subjects using the spherical morphing procedure in FreeSurfer (Fischl et al 1999b).

### 2.2 Forward models

The measured magnetic field **b** at the *N*_c_ sensors 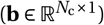 is related to the source space 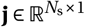 by

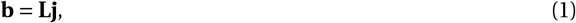

where **L** is the gain matrix 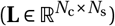 describing the sensitivity of each sensor to a unit source at each location in the source space. Each column *i* of **L** = [**t**_1_, **t**_2_, … **t**_*N*_s__] represents the topography **t***_i_* of source *i* (how the sensors see that source), while each row *j* represents the sensitivity of sensor *j* to all sources. Gain matrices were computed for each subject and sensor array using the linear Galerkin boundary element method (BEM) with an isolated source approach (ISA) as described by Stenroos and Sarvas (2012). The conductivities of the brain, skull and scalp compartments were set to 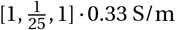, respectively.

### 2.3 Sensor models

The response of each OPM sensor was modelled by a set of eight discrete integration points. These were uniformly distributed within a cube-shaped sensitive volume with a side-length of 3 mm (see Table 1), representing the actual sensitive volume of the sensor. The output of each sensor was computed as the weighted sum of a single magnetic field component over the integration points. SQUID sensors comprised magnetometers (a square pick-up loop with 25.8-mm side length) and planar gradiometers (26.4-mm longer side length, 16.8-mm baseline) as in the Elekta Neuromag^®^ MEG systems (Elekta Oy, Helsinki, Finland); these sensors were modelled as in the MNE software (Table 1). SQUID and OPM magnetometers measured the *z*-component of the magnetic field in the local coordinates specified in Table 1, while the SQUID gradiometers measured a tangential derivative of the *z*-component.

**Table 1.**
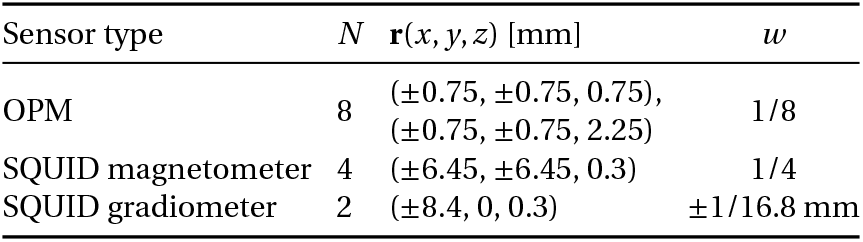
Sensor models. *N* is the number of integration points, **r** are their positions in a local coordinate system, and *w* are their respective weights.

### 2.4 Sensor arrays

OPM sensor arrays were constructed using a subdivision approach in which a spherical surface was divided into horizontal levels. Each level was populated with sensors using equal angular spacing, except in the most anterior part of the head, for which smaller angular spacing was used to compensate for the non-spherical shape of the head. For the lowest levels, a gap was left for the face. This generalised sensor array was then projected onto the scalp surfaces of individual subjects.

The number of sensors per level and the inter-level distance was determined empirically so that the sensor array was maximally dense while still physically feasible for all 10 subjects. The feasibility was determined on the basis of a hypothetical sensor housing with a footprint of 20 mm × 20 mm on the scalp. This criterion resulted in a sensor array consisting of 184 OPMs (Fig. 1).

**Fig. 1:**
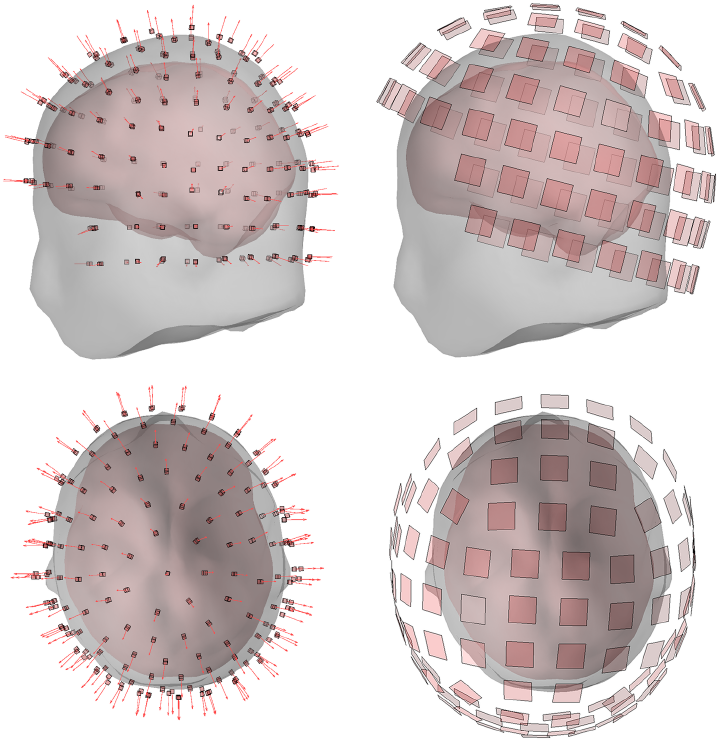
OPM (left) and SQUID (right) sensor arrays for one subject, showing the OPM sensitive volumes and SQUID pick-up loops both from the side and top-down. The red arrows represent the OPM sensitive axes.

The distance from the closest edge of the sensitive volume of the sensors to the scalp was set to 4.5 mm to accommodate a thermal shield and the sensor housing. The sensors were oriented to measure the normal component of the magnetic field with respect to the scalp, as this field component provides most information (Iivanainen et al 2017).

From the original sensor array for each subject, mis-coregistered sensor arrays with sensor-wise random position and orientation errors were constructed. The position of each sensor in a mis-coregistered array was sampled from a uniform distribution within a sphere centred at the true sensor position. The radius of the sphere represented the uncertainty in the measured position of the sensors. Radii were chosen so that the RMS position error was 2, 4 or 6 mm (sphere radii of 2.6, 5.2 and 7.7 mm).

Furthermore, arrays with sensor-wise random orientation error were also constructed. In these arrays, each sensor was tilted from its true axis in a random direction. Error angles were sampled from a uniform distribution such that the RMS orientation error was 5°, 10° and 15° (corresponding to maximum orientation errors of 8.7°, 17.3° and 26.0°). Finally, to approximate more realistic scenarios, sensor arrays with both position and orientation error were constructed; these included arrays with 2-mm and 5°, 4-mm and 10° as well as 6-mm and 15° RMS errors.

To maintain the accuracy of the forward computation, samples were rejected if any of the integration points were less than 2 mm from the scalp surface or below it.

For each simulated error level, 50 different mis-coregistered sensor arrays were constructed per subject, resulting in a grand total of 500 mis-coregistered sensor arrays per scenario.

In addition to the OPM-based sensor arrays, a 306-channel (102 magnetometers, 204 planar gradiometers) SQUID-based Elekta Neuromag^®^ MEG sensor array with the sensor-array position based on experimental MEG measurements was obtained for all subjects. These arrays, which did not include any coregistration errors, were used as the baseline in the comparisons.

We also conducted an additional experiment in which we investigated how the number of sensors in an array affects the robustness to coregistration errors. To this end, we constructed a whole-scalp OPM array consisting of 102 magnetometers by projecting the positions of the SQUID magnetometers in the Elekta Neuromag MEG system onto the scalp. The same 4.5-mm stand-off distance and OPM models as for the 184-OPM array were used. We then constructed erroneous arrays with 6-mm and 15° RMS sensor position and orientation error in the same manner as for the 184-OPM array.

## 3 Metrics and computation

Several measures were used to quantify the effect that mis-coregistration has both on the gain matrices (i.e. the forward models) and on the source estimates calculated using these gain matrices (i.e. the inverse models). All metrics were calculated individually for each subject and each sensor array of that subject. Metrics were thereafter morphed to the average brain of the subjects and averaged. To keep the results informative and to avoid bias due to sources that are practically invisible in MEG, sources were split into deep and shallow ones (Fig. 2). Sources closer than 30 mm from the inner surface of the skull were considered shallow while all others deep, from which MEG signal is less likely to originate.

**Fig. 2:**
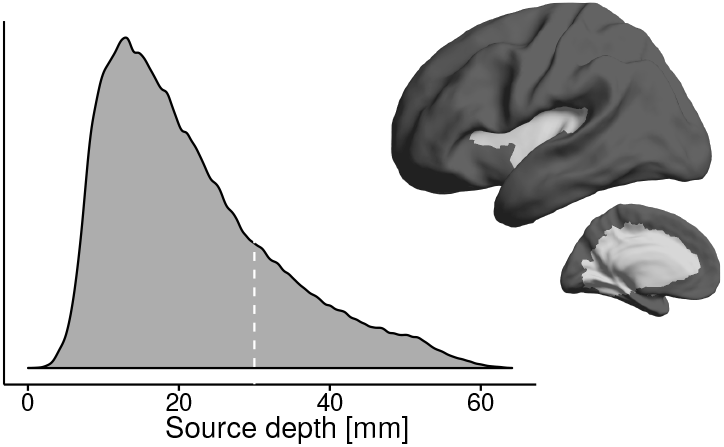
The depth of the cortical sources as measured from the inner surface of the skull (data pooled across subjects): Distribution as a density plot (left). The white dotted line represents the 30-mm threshold at which sources were split into shallow and deep. Mean source depth (right) thresholded to show which areas are classified as deep (light gray) and shallow (dark gray).

### 3.1 Forward metrics

Relative error (RE) is a general difference measure that is sensitive to changes in both amplitude and shape in the topographies. The relative error for the topography of a source is

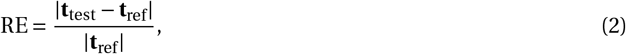

where **t**_ref_ and **t**_test_ are the reference and test topographies of the source, and |·| is the *l*2-norm. **t**_ref_ comes from the gain matrix of the original unperturbed sensor array, while **t**_test_ is from the gain matrix of a mis-coregistered sensor array.

To further investigate the errors in the shape of topographies due to mis-coregistration, the correlation coefficient (CC; Haueisen et al 1997; Stenroos and Sarvas 2012) between the topographies in reference and erroneous sensor arrays were calculated for all individual sources. Unlike RE, this metric is insensitive to amplitude differences. CC is expressed as

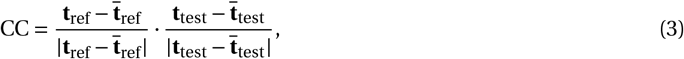

where · is the dot product, and 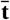 denotes the mean of **t** across all sources.

### 3.2 Inverse metrics

#### Minimum-norm estimation

Minimum-l2-norm estimation (MNE) is a commonly used source estimation procedure (Dale and Sereno 1993; Hämäläinen and Ilmoniemi 1994) in which the minimisation of the *l*2-norm of the source estimate and the fidelity of the reconstruction of the measured signal are balanced. The source estimate 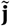 that satisfies the regularised minimum-*l*2-norm criterion is given by

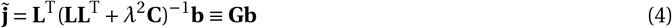

in which *λ*^2^ is the regularisation parameter, **C** is the noise covariance matrix, and **G** is the resulting inverse operator. We used a diagonal noise covariance matrix **C**, whose diagonal elements were proportional to the noise variances of the sensors. The numeric ratio between SQUID magnetometer and planar gradiometer noise variances was 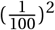 (correspondingto 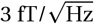 and 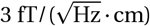), respectively).

We set the regularisation parameter as suggested by Lin and colleagues (2006):

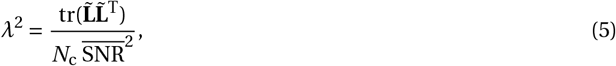

where 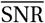 is the assumed mean (amplitude) signal-to-noise ratio, *N*_c_ is the number of sensors, tr(·) is the trace operator, and 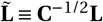 is the whitened gain matrix. We set 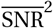 to 9, the default value in MNE software.

We used the minimum-*l*2-norm estimates as the basis for several source localisation metrics. The measures used to investigate the effect of sensor localisation error on the minimum-*l*2-norm source estimates are based on the concept of resolution analysis as applied in earlier literature (Grave De Peralta Menendez et al 1997; Molins et al 2008; Hauk et al 2011; Stenroos and Hauk 2013; Iivanainen et al 2017). In resolution analysis, metrics are derived from the resolution matrix

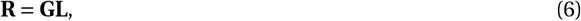

where **G** is the inverse operator, here the MNE operator of Eq. 4. The columns of the resolution matrix **R** consist of point-spread functions (PSFs) 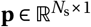, which describe how the activation of each source is distorted by the imaging system, i.e. how the activation of each source is seen in the source estimate. We computed the resolution matrix using the inverse operator based on each mis-coregistered sensor array together with the gain matrix of the original sensor array, thus simulating the effect of mis-coregistration.

We assessed the localisation performance of each sensor array by computing the peak position error (PPE) for all sources, which describes the displacement of the centre-of-mass of the PSF from the actual source position **r**_ref_ (Lin et al 2006; Stenroos and Hauk 2013; Iivanainen et al 2017):

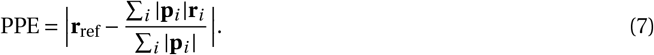

When calculating the PPEs, we thresholded the PSFs by only including the source points at which |**p***_i_*| ≥ 0.5 · **p**_max_, as was done by Stenroos and Hauk (2013).

The spatial spread of the source estimate was characterised by calculating the spatial deviation (SD) of the PSFs (Stenroos and Hauk 2013):

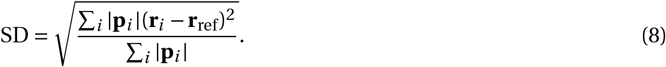

When calculating the SD, we thresholded the PSFs in the same manner as when calculating the PPE.

Finally, as PPE only quantifies the localisation performance of the source estimator, we also computed the correlation between the PSFs of the original and mis-coregistered sensor arrays:

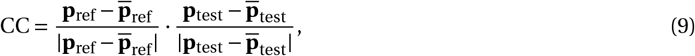

which quantifies the effect of mis-coregistration on the shape of the point-spread functions.

#### Dipole modelling

We implemented a least-squares single-dipole localisation using a simple grid search procedure. The same source space as in the MNE resolution analysis was used as the search space for dipole modelling, but the data were generated using a different source space with non-overlapping discretisation, each source of which was activated separately. To keep the computation time reasonable, the data generation source space was sparser, with an inter-source spacing of 20 mm. The simulated sources were oriented normally to the local cortical surface, but the source orientation was a free parameter in the dipole modelling procedure.

White Gaussian noise was added to the simulated magnetic field to approximate sensor noise. For the SQUID arrays, the noise characteristics of the Elekta Neuromag^®^ MEG system were used: the spectral noise densities for SQUID magnetometers and planar gradiometers were set to 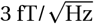 and 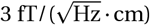, respectively. For the OPMs, three different noise levels were used:

1. 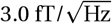, which is equal to that of current SQUID magnetometers.
2. 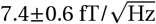, which is the subject-specific break-even noise density at which the SQUID magnetometers and OPMs would have equal SNR (mean±SD across subjects, mean over all cortical sources in the simulation source space).
3. 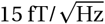, which is demonstrated with current commercial SERF OPMs.

The source amplitude for a given SNR and noise level was determined using the definition of the local SNR for source *i* given by Goldenholz and colleagues (2009):

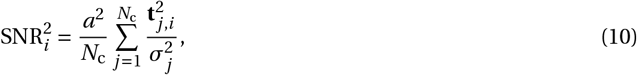

where *a*^2^ is the source variance, *N*_c_ is the number of sensors, and 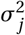 is the noise variance of sensor *j*. In the case of white noise, the noise variance can be expressed in terms of the spectral noise density *n* and bandwidth BW, 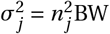. The source variance that produced a mean SNR^2^ of 10 over all sources for the SQUID magnetometer array was used in all simulations, and the bandwidth was set to 40 Hz. This resulted in source amplitudes of 22.1±1.7 nAm across the subjects. To treat both sensor types of the 306-channel SQUID array equally, the gain matrix and data were whitened.

To simulate the effect of mis-coregistration, the simulated magnetic field was computed using the gain matrix of the original sensor array while the gain matrix of a mis-coregistered sensor array was used for the dipole modelling procedure.

We quantified the impact of mis-coregistration using dipole position error (DPE), i.e. the three-dimensional Euclidean distance between the fitted and actual dipoles. In a similar manner, we defined dipole orientation error (DOE) as the difference in orientation between the actual and fitted dipoles. Finally, we compared the goodness-of-fit (GOF), i.e. how well the fitted dipoles explain the original data:

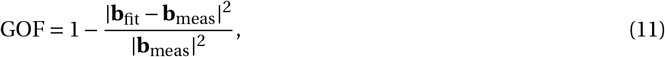

where **b**_meas_ is the measured magnetic field and **b**_fit_ is the magnetic field produced by the fitted dipole.

#### LCMV beamforming

We implemented a linearly constrained minimum-variance (LCMV) beamformer (Van Veen et al 1997; Sekihara et al 2004; Sekihara and Nagarajan 2008). As with the dipole modelling, separate source spaces were used for data generation and source estimation. As the localiser, we used the unit-noise-gain power estimate **Z**^opt^ with optimum orientation:

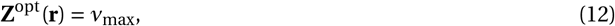

where ν_max_ is the maximum eigenvalue of the matrix

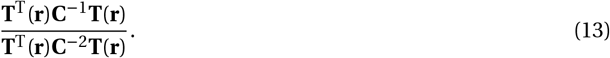

In the above expression, **C** is the data covariance matrix and **T**(**r**) contains the topographies of orthogonal sources at position **r**. The eigenvector corresponding to ν_max_ is the source orientation that produces the maximum power output.

Individual dipolar sources consisting of white Gaussian noise were simulated, and white Gaussian sensor noise was added to the measured magnetic field. The same source amplitudes as in the dipole modelling simulations were used, as well as the noise values corresponding to 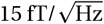 for the OPMs, 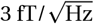 for the SQUID magnetometers and 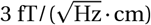 for the SQUID gradiometers. The dataset had a length of 500 samples, and the data covariance matrix **C** was regularized using **C**_reg_ = **C** + *μ* · tr(**C**)/*N*_s_**I** with *μ* = 3. As the simulated sensor noise was white and uncorrelated, this is equivalent to increasing the dataset length while being much cheaper computationally. To treat both sensor types of the 306-channel SQUID array equally, the gain and covariance matrices were whitened before computing the localiser.

Source localisation performance for the LCMV beamformer was quantified by computing the distance between the peak of the **Z**-estimate and the true source location.

## 4 Results

### 4.1 Forward metrics

The relative error of OPM array topographies due to mis-coregistration is shown in Table 2 and Fig. 3. This error is largest in superficial areas, which is to be expected as these areas are closer to the sensors. Similarly, the shapes of the topographies are most affected in superficial areas (Table 2). Unlike the sensor-wise position error, random orientation error is manifested throughout the cortex regardless of source location and depth (Fig. 3 and Table 2). Similar effects can be seen in the shapes of the topographies, although the CCs are not affected to the same degree as the REs. When sensor position and orientation errors are present simultaneously, the effects of these errors add up in a sub-linear manner (Fig. 3).

**Fig. 3:**
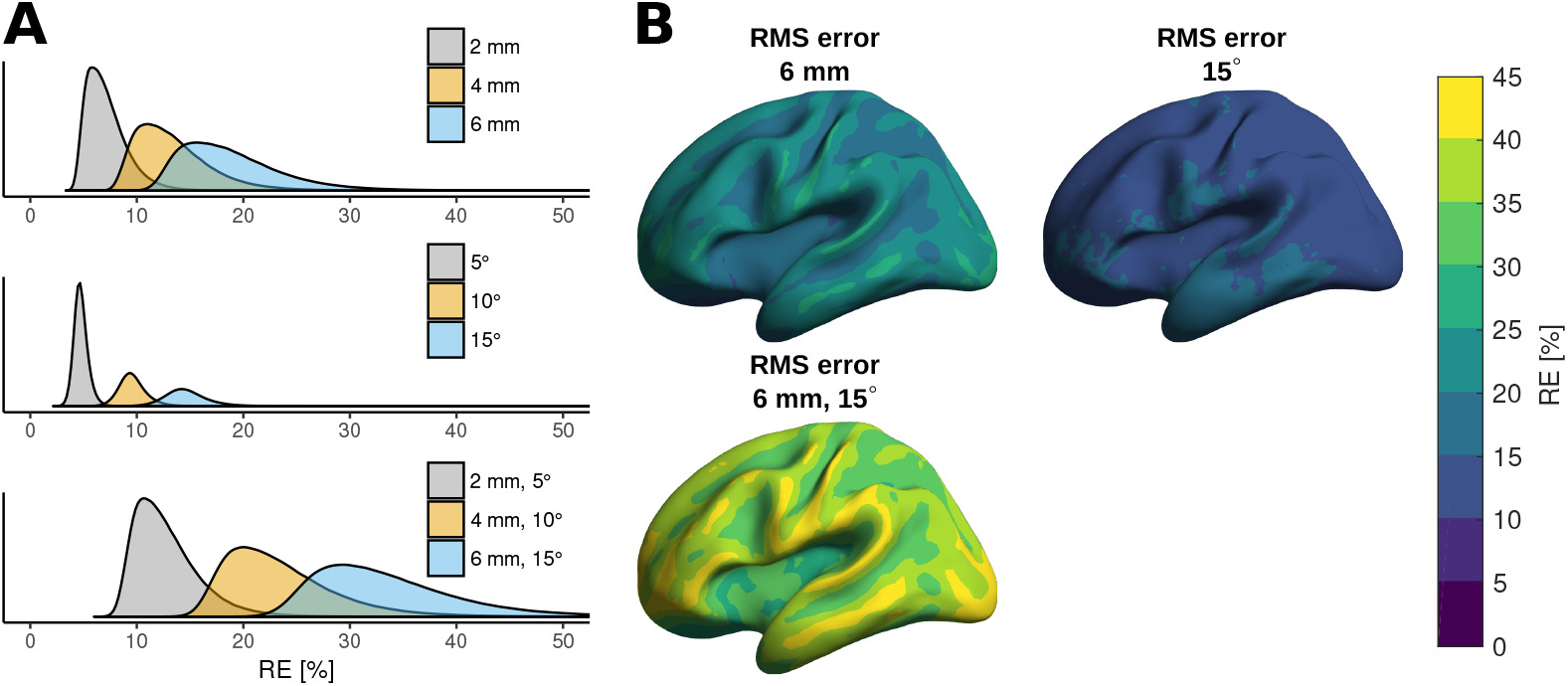
Relative error (RE) of OPM array topographies over all subjects at different levels of sensor position and orientation error. A: Error distributions shown as density plots. B: The mean RE over all subjects.

**Table 2.** Mean relative errors (RE) and correlation coefficients (CC) of source topographies with different levels of sensor-wise position and orientation error.

| RMS sensor-wise error |  | Shallow sources |  | Deep sources |  |
| --- | --- | --- | --- | --- | --- |
| Position [mm] | Orientation [°] | RE [%] | CC [%] | RE [%] | CC [%] |
| 2 | 0 | 7.5 | 99.7 | 5.6 | 99.8 |
| 4 | 0 | 14.1 | 99.0 | 10.4 | 99.4 |
| 6 | 0 | 20.0 | 98.0 | 14.8 | 98.9 |
| 0 | 5 | 4.8 | 99.9 | 4.7 | 99.9 |
| 0 | 10 | 9.7 | 99.5 | 9.4 | 99.6 |
| 0 | 15 | 14.7 | 98.9 | 14.3 | 99.0 |
| 2 | 5 | 13.4 | 99.6 | 10.3 | 99.7 |
| 4 | 10 | 24.5 | 98.5 | 19.3 | 99.0 |
| 6 | 15 | 34.9 | 96.9 | 28.2 | 97.9 |

### 4.2 Inverse metrics

#### Minimum-norm estimation

Peak position errors (PPEs) and correlation coefficients (CCs) of PSFs are summarised in Table 3. The average PPE over all subjects (Fig. 4 and Table 3) is very large for deep sources, of the order of several centimetres, regardless of the amount of sensor position error. The PPEs for shallow sources are much smaller, but are considerable also when no sensor position error is present. The increase in PPE due to sensor position error is seen both in shallow and deep areas of the cortex, although it is proportionally much larger in shallow areas.

**Fig. 4:**
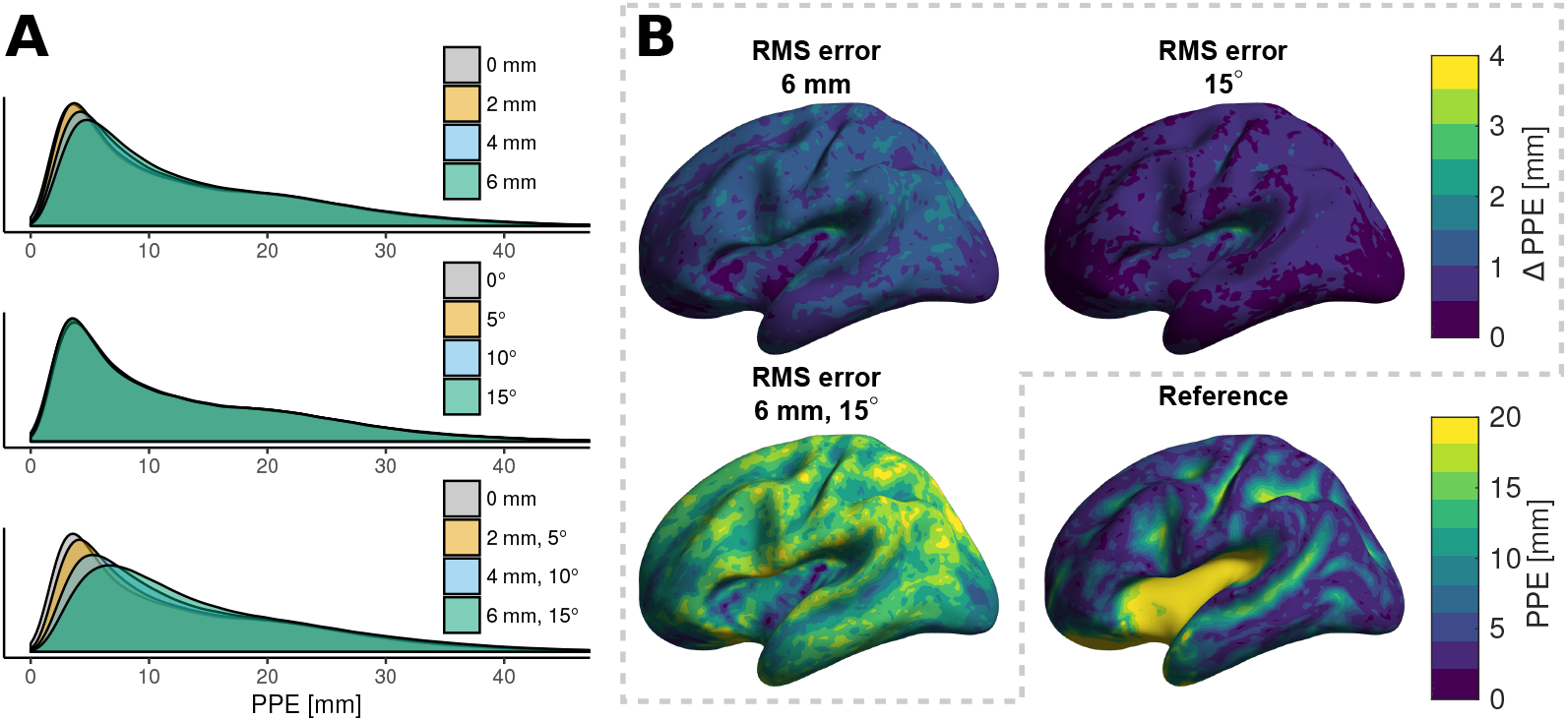
Effects of mis-coregistration on minimum-norm estimation as quantified by the peak position error (PPE) of point-spread functions over all subjects. A: Error distributions as density plots. B: The mean difference in PPE between erroneous and reference sensor arrays.

**Table 3.**
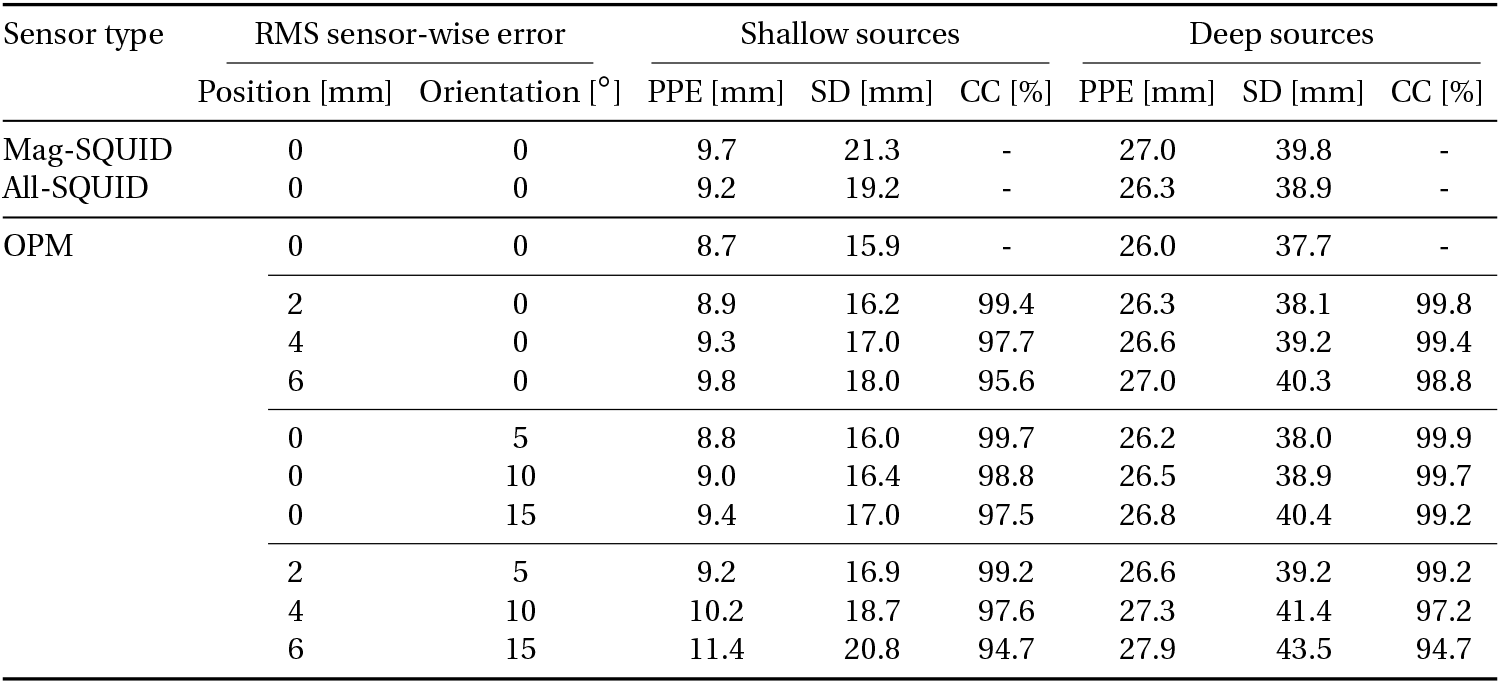
Metrics derived from the point-spread functions computed from the MNE resolution matrix. Mean values of peak position error (PPE), spatial deviation (SD) and correlation coefficients (CC) are listed.

Without mis-coregistration, the PPEs of the OPM array are smaller than those of the reference SQUID arrays, i.e., the 306-channel “All-SQUID” array and the 102-channel magnetometer-only “Mag-SQUID” array (Table 3). The difference diminishes when a small amount of mis-coregistration is present. When the OPM array had both 6-mm and 15° RMS position and orientation errors, the PPEs were larger than those for either (correctly co-registered) SQUID array.

The PSF spread is increased due to mis-coregistration, as quantified by the SD metric (Table. 3). This spreading is most pronounced in areas of high sensitivity. For deep sources, sensor orientation vs. position error causes a proportionately larger SD increase. The SD values of the 102-channel SQUID magnetometer array are much larger than those of the 306-channel SQUID array, which in turn has larger SD values than the OPM array.

#### Dipole modelling

When using the same noise density of 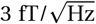 for both SQUID magnetometers and OPMs, the SNR^2^ for the OPMs was 61.6± 10.4 (mean±standard deviation across the subjects). Metrics for this scenario are shown in Table 4. The increased SNR resulted in much larger GOF values and moderately smaller DPEs and DOEs. Even with substantial mis-coregistration, the DPEs for the OPM array remain smaller than those for either SQUID array.

**Table 4.** Metrics related to the accuracy of the dipole fitting procedure at different levels of sensor position and orientation error when both OPMs and SQUID magnetometers had a noise density of 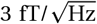 and SQUID gradiometers had a noise density of 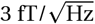. Mean values of position error (DPE), orientation error (DOE) and goodness of fit (GOF) are listed.

| Sensor type | RMS sensor-wise error |  | Shallow sources |  |  | Deep sources |  |  |
| --- | --- | --- | --- | --- | --- | --- | --- | --- |
|  | Position [mm] | Orientation [°] | DPE [mm] | DOE [°] | GOF [%] | DPE [mm] | DOE [°] | GOF [%] |
| Mag-SQUID | 0 | 0 | 12.4 | 50.9 | 81.9 | 26.2 | 56.2 | 56.8 |
| All-SQUID | 0 | 0 | 10.3 | 45.7 | 90.1 | 19.0 | 44.2 | 79.8 |
| OPM | 0 | 0 | 7.3 | 36.1 | 87.3 | 15.8 | 34.6 | 66.1 |
|  | 2 | 0 | 7.6 | 37.0 | 86.9 | 15.8 | 34.6 | 65.9 |
|  | 4 | 0 | 8.0 | 39.2 | 85.9 | 16.0 | 34.9 | 65.4 |
|  | 6 | 0 | 8.5 | 41.7 | 84.5 | 16.3 | 35.4 | 64.8 |
|  | 0 | 5 | 7.4 | 36.4 | 87.1 | 15.8 | 34.6 | 65.9 |
|  | 0 | 10 | 7.5 | 37.1 | 86.7 | 15.8 | 34.7 | 65.5 |
|  | 0 | 15 | 7.7 | 38.2 | 85.8 | 15.9 | 34.8 | 64.8 |
|  | 2 | 5 | 7.6 | 37.2 | 86.7 | 15.8 | 34.6 | 65.7 |
|  | 4 | 10 | 8.1 | 39.8 | 85.2 | 16.0 | 35.0 | 64.9 |
|  | 6 | 15 | 8.7 | 42.7 | 83.0 | 16.4 | 35.8 | 63.5 |

When the noise density of the OPMs was set so that the SNR^2^ were equal to that of the SQUID magnetometers, GOF values were roughly equal for OPM and SQUID magnetometer arrays (Table A1). GOF values of the 306-channel SQUID array were substantially higher. In spite of this, for the shallow sources, the DPEs and DOEs for the OPM array were smaller than those for either SQUID array at all tested levels of mis-coregistration.

Fig. 5 and Table 5 show the dipole modelling results for the simulation scenario where the OPM noise density was set to 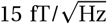, resulting in the OPM SNR^2^ of 2.47±0.42 (Eq. 10, mean±standard deviation across the subjects). The large difference in the SNRs between the SQUID and OPM sensor arrays is clearly seen in the GOF values. In spite of this large difference in SNRs, the dipole localisation accuracy of the OPM sensor array is approximately on par with the 306-channel SQUID array when no mis-coregistration is present. Furthermore, the OPM array has superior localisation accuracy compared to the 102-channel SQUID magnetometer array, even with substantial mis-coregistration.

**Fig. 5:**
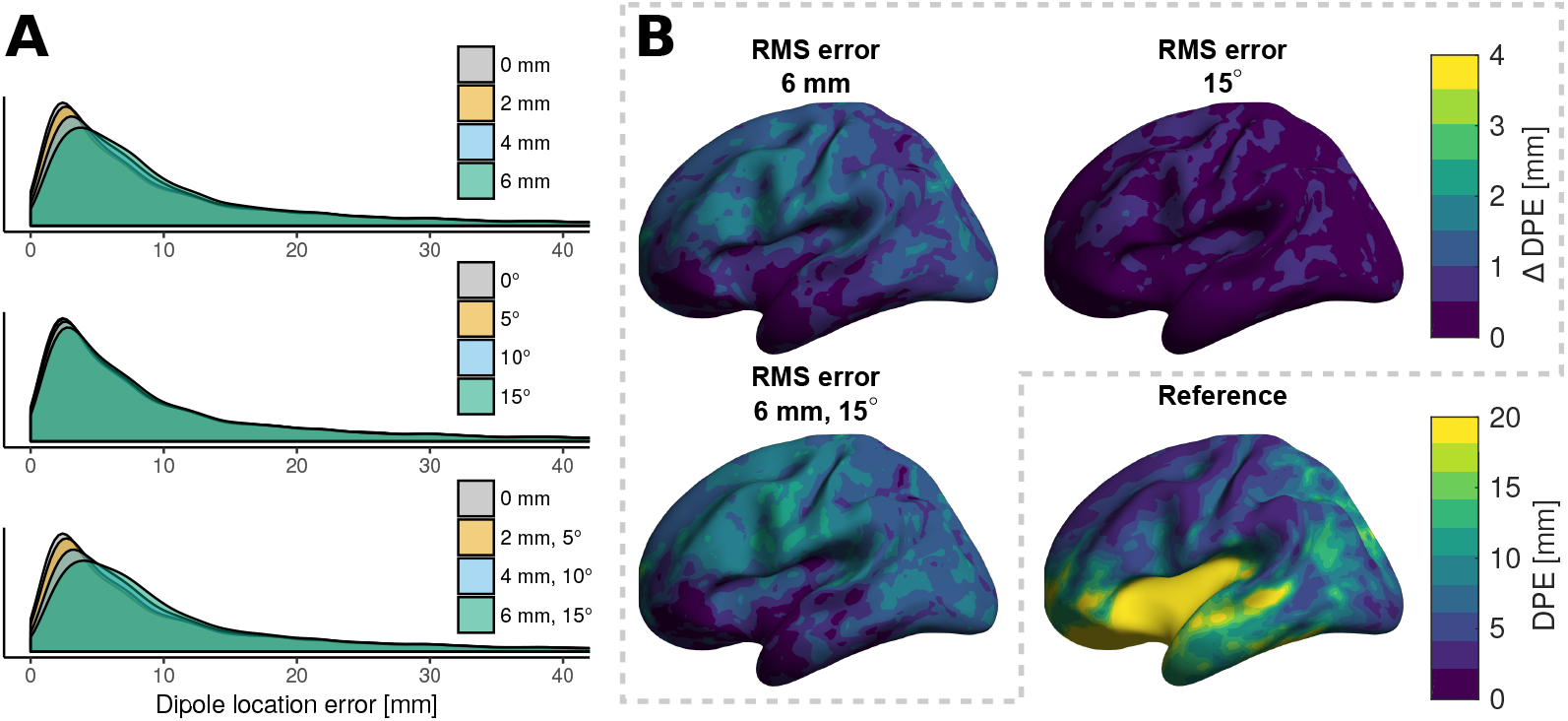
Effects of mis-coregistration on dipole modelling as quantified by the dipole position error (DPE), when OPMs had a noise density of 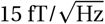. A: Distributions shown as density plots. B: The mean difference in DPE between erroneous and reference sensor arrays.

**Table 5.** Metrics related to the accuracy of the dipole fitting procedure at different levels of sensor position and orientation error when OPMs had a noise density of 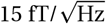, SQUID magnetometers had a noise density of 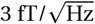 and SQUID gradiometers had a noise density of 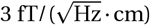. Mean values of position error (DPE), orientation error (DOE) and goodness of fit (GOF) are listed.

| Sensor type | RMS sensor-wise error |  | Shallow sources |  |  | Deep sources |  |  |
| --- | --- | --- | --- | --- | --- | --- | --- | --- |
|  | Position [mm] | Orientation [°] | DPE [mm] | DOE [°] | GOF [%] | DPE [mm] | DOE [°] | GOF [%] |
| Mag-SQUID | 0 | 0 | 12.4 | 50.9 | 81.9 | 26.2 | 56.2 | 56.8 |
| All-SQUID | 0 | 0 | 10.3 | 45.7 | 90.1 | 19.0 | 44.2 | 79.8 |
| OPM | 0 | 0 | 9.7 | 44.0 | 58.9 | 30.9 | 56.5 | 23.3 |
|  | 2 | 0 | 9.9 | 44.5 | 58.7 | 31.0 | 56.6 | 23.3 |
|  | 4 | 0 | 10.3 | 46.0 | 58.0 | 31.2 | 57.0 | 23.1 |
|  | 6 | 0 | 10.8 | 48.0 | 57.0 | 31.6 | 57.5 | 23.0 |
|  | 0 | 5 | 9.7 | 46.2 | 59.6 | 30.9 | 59.6 | 23.3 |
|  | 0 | 10 | 9.9 | 46.7 | 59.3 | 31.0 | 59.7 | 23.0 |
|  | 0 | 15 | 10.1 | 47.7 | 58.8 | 31.2 | 60.0 | 22.6 |
|  | 2 | 5 | 9.9 | 44.6 | 58.6 | 31.0 | 56.7 | 23.5 |
|  | 4 | 10 | 10.3 | 46.7 | 57.6 | 31.2 | 57.1 | 23.0 |
|  | 6 | 15 | 11.0 | 48.7 | 56.0 | 31.8 | 58.0 | 22.6 |

Dipole position errors of < 10 mm are attainable for superficial sources at all sensor position error levels included in the simulations and at all simulated noise levels. The sensor position error has a larger effect on the localisation performance than the orientation error, although the cumulative effect of both error types is even higher. The GOF values are highly affected by source depth and SNR, while mis-coregistration only has a small effect on them.

#### Beamforming

The results for the LCMV beamforming simulations are summarised in Table 4 and Fig. 6. As for dipole modelling, errors of < 10 mm are attainable for shallow sources at all sensor position error levels included in the simulations. Sensor position error has a larger effect on the localisation performance than the orientation error, although the cumulative effect of both error types is even higher. Furthermore, the OPM array has superior localisation accuracy compared to either SQUID array, even with substantial mis-coregistration. The SQUID array including the gradiometers performed slightly worse than the array consisting only of magnetometers, regardless of source depth.

**Fig. 6:**
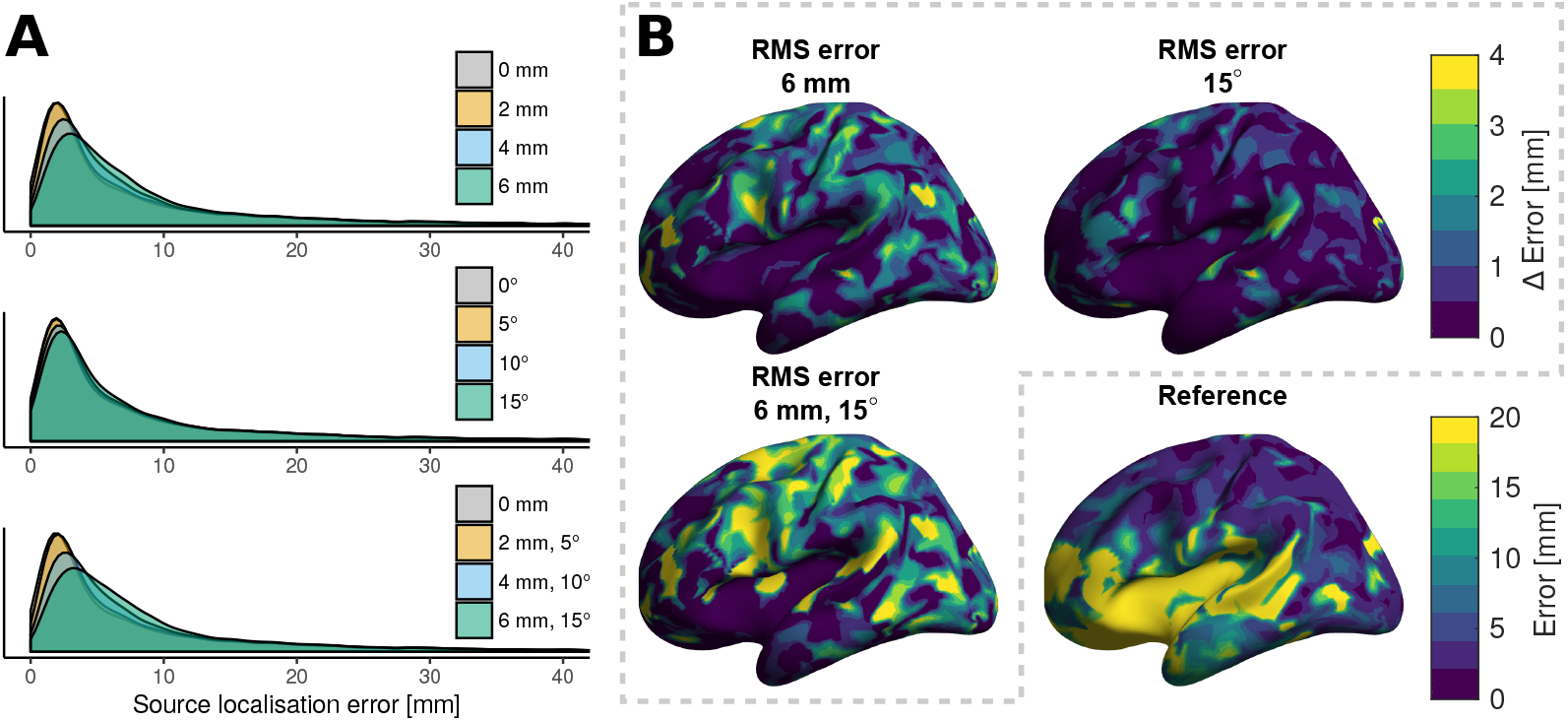
Effects of mis-coregistration on LCMVbeamforming as quantified by the distance between the true source location and the maxima of the beamformer **Z**-estimate. A: Distributions shown as density plots. B: The mean difference in source localisation error compared to the reference array over all subjects at the highest error levels.

### 4.3 Effects of sensor array density

We ran an additional simulation for a 102-sensor OPM array, both without any mis-coregistration and with the maximum level of mis-coregistration used in this study (6-mm RMS position error, 15° RMS orientation error). The amount of forward-model error of the mis-coregistered 102-sensor array was smaller than that of the more dense array (RE for 102- and 184-sensor array: 24.5% and 34.9% for shallow sources, 20.8% and 28.2% for deep sources). However, CCs were similar for both arrays (CC for 102- and 184-sensor array: 97.0% and 96.9% for shallow sources, 97.9% and 97.9% for deep sources). Source estimation results were affected in different ways. For MNE simulations without mis-coregistration, the denser array had slightly better localisation performance than the sparse array (mean PPE for 102- and 184-sensor array: 9.3 mm and 8.7 mm for shallow sources, 26.7 mm and 26.0 mm for deep sources). For dipole modelling, the dense array had higher sensor localisation performance without mis-coregistration (mean DPE for 102- and 184-sensor arrays: 7.7 mm and 7.3 mm for shallow sources, 16.1 mm and 15.8 mm for deep sources). However, when mis-coregistration was present, the MNE results from the sparser array were affected less, to the point that it had superior source localisation accuracy compared to the 184-sensor array (mean PPE for 102- and 184-sensor arrays: 10.8 mm and 11.4 mm for shallow sources, 27.9 mm and 27.9 mm for deep sources). Dipole modelling in the presence of mis-coregistration behaved the opposite way: the sparser array had larger source localisation errors than the denser array (mean DPE for 102- and 184-sensor arrays: 9.5 mm and 8.7 mm for shallow sources, 17.2 mm and 16.4 mm for deep sources). LCMV beamforming performed similarly for both the 102-sensor and 184-sensor arrays, both without (mean source localisation error for 102- and 184-sensor arrays: 7.7 mm and 7.9 for shallow sources, 15.6 mm and 16.2 mm for deep sources) and with mis-coregistration (mean source localisation error for 102- and 184-sensor arrays: 9.5 mm and 9.4 mm for shallow sources, 16.2 mm and 17.0 mm for deep sources).

**Table 6.** Mean error between true source locations and the maxima of the beamformer **Z**-estimate.

| Sensor type | RMS sensor-wise error |  | Shallow sources | Deep sources |
| --- | --- | --- | --- | --- |
|  | Position [mm] | Orientation [°] | Error [mm] | Error [mm] |
| Mag-SQUID | 0 | 0 | 10.7 | 19.6 |
| All-SQUID | 0 | 0 | 11.4 | 21.5 |
| OPM | 0 | 0 | 7.9 | 16.2 |
|  | 2 | 0 | 8.0 | 16.2 |
|  | 4 | 0 | 8.5 | 16.4 |
|  | 6 | 0 | 9.1 | 16.8 |
|  | 0 | 5 | 7.9 | 16.2 |
|  | 0 | 10 | 8.1 | 16.4 |
|  | 0 | 15 | 8.3 | 16.5 |
|  | 2 | 5 | 8.1 | 16.3 |
|  | 4 | 10 | 8.7 | 16.5 |
|  | 6 | 15 | 9.4 | 17.0 |

## 5 Discussion

In this study, we have investigated the effects of mis-coregistration of on-scalp MEG sensors, arranged in non-rigid MEG sensor arrays, on the forward model and source estimation. We found superficial sources to be affected more than deeper ones. We also discovered that the effect of sensor position error was larger than that of the sensor orientation. RMS sensor position errors less than 4 mm increase any of our source estimation error metrics by no more than 8%. Thus, coregistration should not impede the adoption of on-scalp MEG, as long as coregistration methods whose accuracy fulfils our guideline requirement are used.

### 5.1 Simulation methodology

We designed our simulations according to actual, commercially available OPMs (QuSpin Inc., Louisville, CO, USA); the stand-off distance and dimensions of the sensitive volume represented those of these sensors. The hypothetical sensor array was constructed to be maximally dense while still fitting the heads of all 10 adult subjects. The assumed 20 mm × 20 mm scalp footprint of each sensor is a conservative estimate of current OPM casings; QuSpin SERF OPMs have footprints of 13 mm × 19 mm, and smaller yet sufficiently sensitive sensor designs have also been demonstrated (e.g. Sander et al 2012; Alem et al 2014). However, deploying current QuSpin OPMs in an EEG-cap-like sensor array would be challenging due to the 110 mm length of the sensors on the axis normal to the scalp.

The noise density of 15 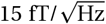 used for the OPMs in the dipole modelling simulations was also based on the QuSpin OPM. For the Elekta Neuromag^®^ SQUID-based MEG system, we assumed 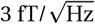 for the magnetometers and 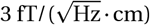 for the gradiometers, which are typical values for this system.

The source estimation methods selected for this study represent very different approaches: minimum-norm estimation assumes an extended source distribution and sets a general, weak prior that aims at reconstructing the relevant part of the measurement with the source distribution that has the minimum *l*2 norm, while dipole modelling uses a very strong assumption of a single focal source. LCMV beamforming, on the other hand, assumes that the data are generated by a small number of temporally uncorrelated sources, and the beamformer scanning function tested the data for dipolar sources. Our simulations were based on single dipoles; the simulated signals thus fulfilled the assumptions behind dipole modeling and beamforming. MNE produces typically rather smeared estimates, making it less accurate in localising focal sources than dipole modelling or beam-forming. Dipole modelling and beamforming are, on the other hand, considered to be more sensitive to errors in the forward model (Stenroos and Hauk 2013; Hillebrand and Barnes 2003). As the simulation setting thus favours dipoles and beamforming over MNE, the localization results from the different source estimation methods should not be compared. Overall, our simulation scenarios were constructed specifically to investigate the effects of mis-coregistration on source estimates, not to compare different source estimation methods.

In the dipole modelling and beamforming simulations, separate and non-overlapping source spaces were used for data generation and source estimation. Thus, any bias that could be caused by using the same source space for simulation and estimation was avoided. This procedure also sets a lower bound for the dipole position error. The closest point in the estimation source space to the data generation source space was on average 1.3±0.6 mm (mean±standard deviation). We did not apply different data generation and source estimation source spaces to the MNE simulations, as we judged the prior of the minimum-norm model to be so different from the simulated single dipoles that the estimation grid does not play a major role when the analysis is done using distributed metrics.

The parameters used for the LCMV beamforming simulations were chosen as to attain a scenario that specifically characterises the effect of mis-coregistration on the source estimate. In particular, we chose a very long effective sampling time. Beamformers are known to react differently to forward model error, or mismatch, than other source estimation methods: In the presence of forward model errors such as mis-coregistration, high input SNRs may be detrimental to source estimation performance. This is well known both within the field of neuroimaging (Hillebrand and Barnes 2003; Dalal et al 2014) and in other fields utilising beamformers (e.g. Cox 1973).

When conducting the simulations, the possibility of intersecting or below-scalp sensor housings was not taken into account as this situation can happen in reality if errors are present in sensor position or orientation measurements. However, such a “sanity check” should be implemented in real coregistration procedures to ensure that the measured sensor positions are reasonable. In particular, if the sensitive volume intersects the head surface, forward modelling may produce large errors. In this study, we rejected sensor locations for which any integration point was closer than 2 mm to the scalp.

### 5.2 Effects of sensor position and orientation errors

Generally, random sensor orientation error produces less adverse effects than position error both on the forward models and source estimation results. Unlike the position error, orientation error affects the forward model throughout the source space regardless of source position or depth. Additionally, sensor orientation errors mostly affected the RE rather than CC values, suggesting that orientation errors primarily alter amplitudes rather than the shapes of topographies. This phenomenon did not directly translate to the source estimation methods included in this study; sensor orientation error mostly affected shallow sources rather than having a global effect.

The localisation performance of MNE for the simulated focal sources is modest with both the OPM and SQUID arrays as demonstrated by the PPE metric, which was never below 5 mm, even without any mis-coregistration. Regardless of the localisation accuracy of MNE, on-scalp MEG systems possess higher spatial resolution than conventional SQUID-based MEG systems, as seen as lower SD values in the current work. Similar findings were also reported by Iivanainen and colleagues (2017).

The higher source localisation performance of the OPM sensor array in comparison to the SQUID arrays was also manifested in the dipole modelling and beamforming simulations; the source localisation error was consistently smaller for OPM arrays even with substantial sensor position error, regardless of the much lower SNR of the OPM arrays. When the OPM noise density was set equal to that of current SQUID magnetometers in the dipole modelling simulations, the SNR of the OPMs was very high, providing a modest improvement in dipole localisation accuracy.

LCMV beamforming results were similar to those attained by dipole modelling, although beamforming had slightly superior source localisation performance (7.9 vs 9.7 mm without mis-coregistration). The performance difference can be explained by the amount of time-domain information used: The LCMV source estimate is based on a data covariance matrix from multiple samples while the other estimators used only single samples.

Interestingly, the full SQUID array with both planar gradiometers and magnetometers performed worse in LCMV beamforming than the SQUID array consisting only of magnetometers. In fact, utilising only the SQUID planar gradiometers in the beamformer resulted in worse performance (mean source localisation error for shallow and deep sources was 18.2 and 19.1 mm, respectively). This counterintuitive result is due to the absence of brain noise in the simulations, which causes the magnetometers to have much larger SNRs than the gradiometers. It would seem that the lower SNRs of the gradiometers partially corrupt the data covariance estimate, worsening the source localisation performance of the SQUID array with both sensor types.

The relative effect of mis-coregistration on source localisation accuracy was most pronounced for MNE, as shown by the PPE metric. Dipole modelling was less susceptible to mis-coregistration when OPMs had a noise density of 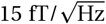, however, when the SNR was increased by changing their noise density to 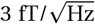, the relative effect of mis-coregistration was increased as it was no longer masked by noise. LCMV beamforming was more sensitive to mis-coregistration than dipole modelling at equal SNRs, but not as sensitive as MNE.

### 5.3 Requirements for sensor localisation accuracy

Since the SNR of MEG diminishes with increasing source depth, shallow neocortical sources typically dominate MEG signals. Therefore, in the following we focus on the results for shallow sources. According to our results, an RMS sensor position error of 4 mm increases the source localisation error by 7% for MNE (change in PPE), 10% for dipole modelling (change in DPE, OPM noise density: 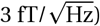) and 8% for LCMV beamforming compared to error-free coregistration. We also included simulations scenarios in which both position and orientation errors were present. For example, combined 2-mm and 5° RMS errors had < 6% effect on sensor localisation accuracy for any of the tested source estimation methods. However, RMS errors of 4 mm and 10° increased source localisation errors by 17%, 11% and 10% for MNE, dipole modelling and beamforming, respectively. For MNE, the spatial spread of the estimate also increases with mis-coregistration: for example, RMS errors of 4 mm and 10° increases by 18%.

Ultimately, it is up to the user to decide how accurate source estimates are needed for their specific research question and methodology, and thus by extension how large sensor position and orientation errors can be tolerated. Nevertheless, on the basis of our simulation results, we propose 4-mm sensor position and 10° sensor orientation RMS errors as a general guideline for the maximum acceptable mis-coregistration for large on-scalp MEG systems. When coregistration errors are smaller than these requirements, the on-scalp MEG system performed at or above the level of conventional SQUID-based MEG systems. With larger coregistration errors, the advantage of on-scalp MEG may be lost.

These guideline requirements are appropriate for the inverse modelling techniques presented in this study. Compared to those techniques, other distributed source estimates which pose strong priors, typically sparsity of the estimate, may be more sensitive to mis-coregistration. These methods include minimum-*l*1-norm estimation (Matsuura and Okabe 1995; Uutela et al 1999), mixed-norm estimation (Ou et al 2009; Gramfort et al 2012) and techniques utilising multivariate source prelocalisation (Mattout et al 2005).

Interference suppression techniques based on the physics of the measurement, such as the signal-space separation (SSS) method (Taulu and Kajola 2005) and variations thereof, may also have stricter coregistration requirements. For example, Nurminen and colleagues (2008) showed that highly accurate knowledge of array geometry was necessary for both SQUID magnetometer and axial gradiometer arrays to reach high suppression factors against external magnetic field sources when using SSS. Additionally, they reported that these accuracy requirements are likely to be even stricter for sensor arrays capable of measuring higher spatial frequencies, such as on-scalp MEG systems.

Furthermore, the number of sensors may affect the robustness of source estimation results in the presence of mis-coregistration. We constructed a sparser 102-sensor OPM array and compared it to the 184-sensor OPM array. When the sensor density is increased, co-registration errors of the same absolute level become larger in relation to the inter-sensor distance, which was reflected in the larger forward-model relative errors (RE) of the dense array. Since the topography correlation coefficients (CC) were similar, the larger REs of the dense array stem primarily from relative sensitivity changes without changes in topography shape.

Due their differing sensitivities to RE, the robustness of source estimation methods could depend differently on the channel count. We found that for MNE the sparser array had slightly worse source localisation accuracy when no coregistration errors were present, but this sparser array was also more robust to mis-coregistration. For dipole modelling, this was not the case; in the presence of mis-coregistration, the additional sensors in the 184-sensor array provided more error tolerance. LCMV beamforming was not affected by the number of sensors to the same extent; source localisation accuracy was similar for both the 102-sensor and 184-sensor arrays, regardless of mis-coregistration.

The results of this study can to some extent be generalised to other novel sensor types than OPMs, as long as the sensors are deployed in non-rigid arrays. For example, coregistration requirements for arrays comprising high-T_C_ SQUIDs are likely similar although it should be taken into account that these sensors have a planar sensitive area while OPMs have a sensitive volume.

### 5.4 Sensor localisation methods

In light of our results, most of the sensor localisation methods that have been applied in MEG and EEG seem to fulfil our guideline requirements; these methods yield < 4-mm RMS errors in a head-sized volume when used with care, and should thus guarantee source estimation performance at least on the level of SQUID-based MEG. For example, the Polhemus^®^ 3D electromagnetic digitiser system (Polhemus Inc., Colchester, VT, USA) has a reported accuracy of 3–7 mm (e.g. Koessler et al 2010; Baysal and Şengül 2010; Dalal et al 2014). Methods more accurate than the Polhemus^®^ system exist, especially the recently-developed optical ones such as photogrammetry (e.g. Bauer et al 2000; Russell et al 2005) and structured-light scanning techniques (e.g. Koessler et al 2011; Ettl et al 2013; Hironaga et al 2014). However, other factors such as cost, ease of use, coverage, and speed may be the decisive factors in choosing the method. For example, the Polhemus ^®^ system requires the operator to move a digitiser stylus to every sensor, making coregistration more laborious and error-prone. This holds true especially if one needs to digitise the sensor orientation as well, requiring one to digitise at least two points along the sensor normal. To its benefit, the Polhemus^®^ system is proven to work reliably and is widely used for coregistration for both EEG and MEG. Optical surface mapping methods, on the other hand, have the potential to be fast (Koessler et al 2011) or even instantaneous (Bauer et al 2000) depending on the implementation. In contrast to e.g. the Polhemus system, sensor orientation can be determined from optical scan surface data without any additional measurement steps. When using optical methods, one needs to identify the individual sensors non-ambiguously. In spite of this complication, optical surface mapping methods seem a very promising solution, as they can collect large amounts of surface data, including the shape of the head and face, very quickly (Koessler et al 2011). However, line-of-sight issues may hinder the use of optical coregistration methods depending on how the sensors are mechanically supported. Additionally, some coregistration methods may not be suited to continuous use during measurements, which limits their practical benefits.

In SQUID-based MEG, the Polhemus^®^ system is the most widely used coregistration method, although some developments using the methods described above have taken place; optical scanning systems have been applied to MEG (Bardouille et al 2012; Hironaga et al 2014; Murthy et al 2014), as well as individualised head casts that snugly fit between the head of the subject and the helmet-shaped cavity of the dewar (Troebinger et al 2014; Meyer et al 2017). Similar efforts have been devoted to OPM-based on-scalp MEG by Boto and colleagues (2017), who used a head cast for both coregistration purposes and to physically support the sensors. When the sensors are mounted in a fixed geometry, such as a head cast, the coregistration problem will be similar to that of current SQUID-based MEG, where systematic shifts of the array are the dominant error sources.

### 5.5 Sources of model error

Sensor localisation is just one part of the coregistration process and thus not the only source of coregistration error. Once the position and orientation of the sensors and some head surface points (e.g. fiducials) are known in a common coordinate system, the surface points are typically fitted to MR-images. A variety of methods for this fitting procedure exist, e.g. adaptations of the Levenberg–Marquardt algorithm (Kozinska et al 1997, 2001) and the ICP algorithm (Besl and McKay 1992). Coregistration procedures using solely the standard anatomical landmarks (the nasion and the preaurical points; Jasper 1958) are considered to be less accurate than those that additionally use dense head surface data (Whalen et al 2008). In the current study, mis-coregistration was attributed exclusively to errors in the measurement of sensor position and orientation.

### 5.6 Prospects

To date, several research groups as well as commercial entities have developed OPMs that can be and have been applied to MEG both in humans (e.g. Shah and Wakai 2013; Johnson et al 2013) and animals (Alem et al 2014). The sensors and experimental set-ups vary from single-channel measurements using physically very large magnetometers (e.g. Xia et al 2006; Kamada et al 2015) to compact sensors (e.g. Shah and Wakai 2013; Knappe et al 2014) that could feasibly be deployed in a sensor array covering the entire scalp. Even though single sensors theoretically suitable for a high-density whole-head array have been developed, no such system has been demonstrated yet. Remaining challenges, apart from coregistration, include improvements in magnetic shielding and reduction of sensor cross-talk as well as lowering total system cost. An additional practical challenge for OPM-based MEG is sturdy but adjustable mechanical support for a high-density OPM sensor array. Using a cap as in EEG might be viable, but it remains to be seen if such a solution provides enough mechanical support for OPMs, which are substantially heavier and larger than EEG electrodes and are sensitive to changes in their orientation.

### 5.7 Limitations

Some factors that may cause issues related to coregistration were not taken into account in the present work. We did not include sensor calibration errors, which may be a significant source of error for OPM-based source estimation, in our simulations. However, the effect of sensor-wise calibration error should be similar to that of sensor-wise coregistration error, as both error types will distort lead fields in a sensor-wise manner. Furthermore, we concentrated exclusively on the effects of sensor-wise coregistration errors. Systematic coregistration errors, as with conventional MEG and EEG, may also occur, and their effects were not examined in the current work.

Although we touch upon the effects of on-scalp MEG sensor array density in this work, that is not our primary focus. For more extensive coverage of this issue, see e.g. Boto et al (2016) and Riaz et al (2017). We did not investigate any partial-coverage sensor arrays, as the characteristics of partial-coverage arrays (and their source estimation performance) must be considered on a case-to-case basis, with factors such as scalp coverage and sampling density being decided based on the expected signal sources and the specific research question.

## 6 Conclusions

In this work, we investigated the effect of errors in sensor positions and orientations in on-scalp MEG arrays. We found that position error introduces larger errors in the forward model and hampers source estimation performance more than the orientation error does. Based on our results, we propose < 4-mm RMS position and < 10° RMS orientation error levels as a general-purpose requirement for source estimation in on-scalp MEG. Current coregistration methods used in both EEG and MEG generally fulfil these requirements; thus, coregistration should not pose a large problem to the adoption of on-scalp MEG. Yet, there is a need for a faster, more reliable and practical method for determining on-scalp MEG sensor positions and orientations.

## Acknowledgements

The authors thank the Advanced Magnetic Imaging centre of Aalto Neuroimaging Infrastructure for the MR images and Aalto University Science-IT for providing computational resources. Research reported in this publication received funding from the European Research Council under ERC Grant Agreement n. 678578 and the National Institute of Neurological Disorders and Stroke of the National Institutes of Health under Award Number R01NS094604. The content is solely the responsibility of the authors and does not necessarily represent the official views of the funding organisations.

## Compliance with Ethical Standards

### Conflict of interest

The authors declare that they have no conflict of interest.

### Ethical approval

For this type of study formal consent is not required.

## Appendix

**Table A1:** Metrics related to the dipole fitting procedure at different levels of sensor position and orientation error when the noise density of OPMs was set so that their SNR was equal to that of SQUID magnetometers. SQUID magnetometers had a noise density of 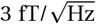 and SQUID gradiometers had a noise density of 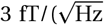. The resulting noise density for the OPMs was 7.4±0.6 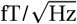. Mean values of dipole position error (DPE), orientation error (DOE) and goodness of fit (GOF) are listed.

| Sensor type | RMS sensor-wise error |  | Shallow sources |  |  | Deep sources |  |  |
| --- | --- | --- | --- | --- | --- | --- | --- | --- |
|  | Position [mm] | Orientation [°] | DPE [mm] | DOE [°] | GOF [%] | DPE [mm] | DOE [°] | GOF [%] |
| Mag-SQUID | 0 | 0 | 12.4 | 50.9 | 81.9 | 26.2 | 56.2 | 56.8 |
| All-SQUID | 0 | 0 | 10.3 | 45.7 | 90.1 | 19.0 | 44.2 | 79.8 |
| OPM | 0 | 0 | 7.9 | 38.2 | 77.7 | 20.1 | 42.0 | 43.6 |
|  | 2 | 0 | 8.0 | 39.0 | 77.4 | 20.1 | 42.0 | 43.5 |
|  | 4 | 0 | 8.5 | 41.1 | 76.5 | 20.3 | 42.5 | 43.2 |
|  | 6 | 0 | 9.0 | 43.3 | 75.2 | 20.7 | 43.1 | 42.8 |
|  | 0 | 5 | 7.9 | 38.4 | 77.3 | 20.1 | 42.1 | 44.3 |
|  | 0 | 10 | 8.1 | 39.1 | 76.1 | 20.2 | 42.2 | 42.2 |
|  | 0 | 15 | 8.2 | 40.1 | 76.4 | 20.3 | 42.3 | 42.8 |
|  | 2 | 5 | 8.1 | 39.2 | 77.2 | 20.2 | 42.1 | 43.4 |
|  | 4 | 10 | 8.6 | 41.6 | 75.9 | 20.4 | 42.5 | 42.9 |
|  | 6 | 15 | 9.2 | 44.3 | 73.9 | 20.9 | 43.5 | 42.1 |

